# The Magic Impacts of Sounds on Consumer’s Brain: How Do Natural Sounds Nudge Green Product Purchase?

**DOI:** 10.1101/2024.02.29.582721

**Authors:** Geying Liang, Yiwen Wang

## Abstract

Despite extensive research on nudging techniques aimed at promoting green consumption, the influence of natural sounds on encouraging environmentally friendly behavior remains relatively understudied. It is valuable to examine how sounds matter in green marketing, as more and more firms are using background sounds in video advertisements. Drawing on the Stimulus-Organism-Response model, we propose that natural sound increase early attentional congruency associated with green products, thereby promoting customers’ purchase of green products. To test on our theory, we conducted an experiment using an event-related potentials (ERP) method and employed a hierarchical drift-diffusion model (HDDM). The results show that compared with unnatural sounds, natural sounds not only increased the purchase rates for green products but also enhanced decision-making efficiency (drift rate) in favor of purchasing green products. Additionally, consumers also exhibited a reduced frontal early P2 wave (150-230ms) in response to green products under natural sounds, indicating that natural sounds increased the early attentional congruency associated with green products. More importantly, the early attentional congruency provides an explanation for how natural sounds nudge the efficiency of purchasing decisions for green products. This study thus contributes to the discovery of a new psychological path that helps deepen our understanding of how natural sounds can nudge green consumption. It also provides actionable implications for green consumption market managers.

## 1. Introduction

The term “green consumption” refers to the deliberate and conscious selection, utilization, and appropriate disposal of goods and services in a manner that is environmentally responsible and promotes sustainable development (Carlson et al., 1993). How to promote a consumer behavior that aligns with this ethical business practices is a highly regarded issue in global business development (García-de-Frutos et al., 2018; Lee et al., 2014; Zheng et al., 2023). For instance, European Union (EU) member states such as the United Kingdom, Denmark, and Sweden provide consumers with information about environmentally friendly products through ecolabeling (Sønderskov & Daugbjerg, 2011). However, a recent survey indicates that EU citizens lack confidence in being able to engage in green consumption in the actual market, with only 46% of Europeans believing that green consumption can be fully implemented (Eurobarometer, 2022). Furthermore, an empirical study conducted in Indonesia has found that even though people hold favorable attitudes towards green consumption, they may not necessarily choose green products in their actual consumption behavior (Arli et al., 2018). A recent meta-analysis has also found that studies conducted in both Western and non-Western countries have explored the phenomenon of positive attitudes towards green consumption not translating into corresponding behavior (Zaremohzzabieh et al., 2021). All of these indications suggest that traditional green consumption propaganda has a limited effect. Therefore, some researchers have found that the nudge approach has been highly effective in promoting green consumption. For example, Lee et al. (2020) found that using videos to draw consumer attention to the natural environment, rather than the knowledge about green products, can nudge their preference for green-labeled fashion products. Therefore, this study proposes the role of natural sounds as a simple and easily processed stimulus that can potentially nudge green consumption.

Natural sounds refer to sounds in nature, including birds (Ratcliffe et al., 2013; Spendrup et al., 2016), waterfalls (Koivisto et al., 2022), rain (Lin et al., 2022), and so on. Many previous studies have shown that sounds have a significant impact on consumer behavior, but most of the focus has been on background music (Biswas, 2019; Michel et al., 2017; North et al., 2003). The earliest study on the impact of natural sounds on consumption, as found in our literature review, was a field experiment conducted by Spendrup et al. (2016) (see Table 1). Subsequently, an increasing number of scholars have explored the influence of natural sounds on consumption. Although previous research has provided some profound insights into the facilitation of consumption by natural sounds, there still exist certain limitations.

**Table 1.** Summary of selected research on the relationship between natural sounds and consumption.

|  | Studies | Sound conditions | Product types | Consumption metrics | Psychological or physiological metrics | Measures | Research designs | Theoretical (or framework) support | Key findings |
| --- | --- | --- | --- | --- | --- | --- | --- | --- | --- |
| 1 | Spendrup et al. (2016) | No sounds, natural sounds(bird) | Organic foods, climate friendly foods, control | Willingness to buy (WTB) | Connectedness to nature | Self-reported | Field experiment | Environmental psychology model | 1.The study revealed a positive and direct impact of natural sounds on the purchase intentions of male customers who initially had low intentions to buy organic foods.<br>2. Sustainability information positively moderated the WTB both organic and climate-friendly foods in men with high connectedness to nature. |
| 2 | Lin et al. (2019) | Park sounds, food court sounds, fast food restaurant sounds, café sounds, and bar sounds | Chocolate gelato | Flavour trajectories | Affective responses | Self-reported | Lab experiment | Emotional mediation theory | 1. Park and café sounds increased the perception of sweetness during the early mastication period.<br>2. The park sound received a significantly higher valence rating, while the bar sound obtained a significantly higher arousal rating.<br>3. The pleasant park sound correlated positively with the perceived sweetness of the gelato. |
| 3 | Xu et al. (2019) | Café sounds, café-bird sounds, café-forest sounds, café-machine sounds | Chocolate, ice-cream | Flavour trajectories | Emotional responses, heart rate, skin conductance, respiration, blood volume pulsation | Self-reported , electrophysiological measures | Lab experiment | Circumplex model of emotion | 1. The café-forest soundscape intensified the perception of sweetness and creaminess at the start of consumption.<br>2. The café-bird soundscape enhanced the creaminess during consumption.<br>3. Both the café-forest and café-bird soundscapes significantly increased positive emotions.<br>4. Exposure to the café-forest soundscape was associated with increased perceptions of sweetness and positive emotions in relation to the ice cream, potentially leading to a relaxed state, as indicated by an increase in blood volume pulse amplitude. |
| 4 | Peng-Li et al. (2021) | Healthy sounds (ocean waves and seagulls), unhealthy sounds (traffic noise) | Healthy foods, unhealthy foods | Food choice | Visual attention (Fixation duration, count, revisit count) | Eye-tracking food choice paradigm | Lab experiment | Dual-system model, gaze bias theory | 1. The healthy soundtrack increased the selection of healthy foods.<br>2. The healthy soundtrack heightened fixations on healthy food items.<br>3. Sound partially mediated food choices through fixation duration, count, and revisit count. |
| 5 | Peng-Li, et al. (2022) | Natural sounds (oceans waves), Noise (chattering and tableware noises) | Healthy and savory foods, healthy and sweet foods, unhealthy and savory foods, unhealthy and sweet foods | Explicit liking, explicit wanting, implicit wanting, reaction time | No | Leeds food preference questionnaire (self-reported , paired food choice) | Lab experiment | The model integrates the incentive-sensitization theory | 1. Natural sounds increased explicit liking for healthy foods over unhealthy foods but had no effect on measures of wanting (explicit or implicit).<br>2. The presence of restaurant noise, as opposed to natural sounds, led to quicker response times for both nutritious and less healthy food options. |
| 6 | Lin, et al. (2022a) | Liked music, pleasant sounds (such as birds, waterfalls, rain), liked music with pleasant sound, disliked music, and disliked music with unpleasant sound (such as busy traffic) | Chocolate ice-cream | Flavour trajectories | Emotion | Self-reported | Lab experiment | Valence–arousal–dominance model | Eating ice cream with liked music and pleasant sounds generated more positive emotions than merely listening to pleasant sounds. |
| 7 | Lin et al. (2022b) | Unpleasant sounds (such as traffic, indoor marketplace, inside of airport), pleasant sounds (such as bird, waterfall, rain) | Emotional foods, non-emotional foods | Flavour trajectories | Emotion | Self-reported | Lab experiment | Valence–arousal–dominance model | In chocolate milkshakes, pleasant sounds had a greater effect on positive emotions and sweet perception compared to vegetable ice creams. |
| 8 | Michels and Hamers (2023) | Bird sounds, water sounds, wind sounds | Healthy foods, unhealthy foods | Food craving, snack intake | Negative affect, salivary cortisol, heart rate variability, hunger, influence expectations | Self-reported, electrophysiological measures | Lab experiment | Stress reduction theory, attention restoration theory | 1. Both bird and water sounds were more effective at promoting cortisol recovery and reducing stress compared to wind sounds alone.<br>2. Water sounds had the most pronounced effect on cortisol recovery. |
| 9 | This study | Natural sounds (birds and wind), unnatural sounds (noise of machines) | Green products, non-green products | Purchase rate, decision efficiency (drift rate) | Early attentional congruency (P2) | Hierarchical drift-diffusion model, event-related potential method | Lab experiment | SOR model, attention restoration theory, musical congruity hypothesis | 1. Natural sounds promoted green consumption (purchase rate).<br>2. Natural sounds lead to more frequent and quicker purchase decisions for green products (drift rates).<br>3. The decreased P2 component suggests that natural sounds increase the early attentional congruency.<br>4. Increased attentional congruency is correlated with increased decision efficiency. |

*Note: 1. The text within parentheses provides an explanation of the current word or serves as a measurement indicator for the current word. 2. In our literature review, we only presented the significant differences in indicators or findings related to natural sounds.*

Firstly, upon reviewing the previous studies (see Table 1), we discovered that the majority of them are focused on the influence of natural sounds on food consumption. These consumption metrics encompass the willingness to buy for organic and environmentally friendly foods (Spendrup et al., 2016), the perception of food flavors (Lin et al., 2019; Lin et al., 2022a; 2022b; Xu et al., 2019), the choice between healthy and unhealthy foods (Peng-Li et al., 2021), the liking and wanting of different flavors of healthy and unhealthy foods (Peng-Li et al., 2022), as well as the craving and intake of healthy and unhealthy foods (Michels & Hamers, 2023). However, it remains unknown whether natural sounds would influence the actual purchasing decisions of green products beyond the food category. Furthermore, the binary purchase decision involves efficiently evidence-accumulation process for or against buying a product until a decision threshold is reached (Krajbich et al., 2012). This efficiency not only includes whether consumers purchase green products but also how quickly they make decisions to purchase green products. Without considering the efficiency of the evidence-accumulation process in consumers’ purchase decision-making, marketers will struggle to develop effective long-term green product marketing strategies. Therefore, it is crucial to explore the effect of natural sounds on the efficiency for in consumers’ green product purchase.

Secondly, previous research has found that natural sounds have an impact on various psychological and physiological metrics (see Table 1). These metrics include connectedness to nature (Spendrup et al., 2016), affective or emotional responses (Lin et al., 2019; Xu et al., 2019; Lin et al., 2022a; 2022b; Michels & Hamers, 2023), visual attention (Peng-li et al., 2021), heart rate (Xu et al., 2019), and more. According to attention restoration theory (Kaplan, 1995), natural sounds can mitigate the depletion of people’s attentional resources. Moreover, from an evolutionary perspective, natural sounds (such as bird sounds) may have a rapid and less perceptible effect on consumers (Kellert & Wilson, 1993). This suggests that natural sounds may impact consumers’ early attentional mechanisms. Moreover, the music congruity hypothesis proposes that matching background sounds with the perception of products can increase consumers’ preference for them (Alpert & Alpert, 1990; North et al., 2016). There is a possibility that natural sounds may influence early attentional congruency. Therefore, investigating early attentional congruency is critical for comprehending the impact of natural sounds on green products during the processing of product information.

Finally, prior research has found that natural sounds can influence consumers’ psychological or physiological states, thereby impacting their purchasing intentions (Spendrup et al., 2016; Peng-Li et al., 2021). For example, Peng-Li et al., (2021) discovered that natural sounds of waves and beaches enhanced consumers’ visual attention towards healthy foods, leading to an increased choice for healthy foods. Visual attention is a bottom-up attentional mechanism in consumers, which is influenced by both the environment and task at hand (Atalay et al., 2012; Clement et al., 2015). From a temporal perspective, it remains unclear whether natural sounds influence individuals’ decision-making by impacting consumers’ early attentional congruency. Furthermore, previous research has only identified effects on choice, and it remains unknown whether natural sounds further impact decision efficiency through early attentional congruency. Therefore, we aim to further investigate whether natural sounds may impact the early attentional mechanism involved in processing product information, thus affecting the efficiency of decision-making regarding the purchase of green products.

In summary, based on these research gaps, this study poses the following questions: 1. Does natural sound enhance the efficiency of consumers’ purchase decisions for green products? 2. Does natural sound influence the early attentional congruency involved in processing information about green products among consumers? 3. Is the early attentional congruency influenced by natural sounds a contributing factor to the efficiency of decision-making for green products?

To answer these questions, we have developed a theoretical framework based on the Stimulus-Organism-Response (S-O-R) model (Mehrabian & Russell, 1974; Jacoby, 2002). The S-O-R model proposes that environmental factors can influence consumer responses by impacting their internal factors, thereby affecting their responses. Firstly, we utilize an experimental rule of randomly implementing a single trial of the task (Knutson et al., 2007). Therefore, the decisions made by the participants in this study were incentivized purchasing decisions. To investigate the influence of natural sounds on the efficiency of the evidence accumulation process for making purchasing decisions regarding green products, we employed a hierarchical drift diffusion model (HDDM). Secondly, we used an event-related potentials (ERP) method to capture consumers’ early neural signals related to attentional congruency. Thirdly, we propose to establish a model with the early attentional congruency as a mediator to examine the nudging effect of natural sounds on the efficiency of decision-making for green products.

Our study contributes to the following literatures. Firstly, we explored the impact of natural sounds on the efficiency of consumers’ decision-making in purchasing green products from an evidence accumulation perspective. Secondly, we enhance our understanding of the attentional mechanisms through which natural sounds influence the processing of information about green products from a temporal perspective. Thirdly, we contribute to the application of the S-O-R model by examining how natural sounds enhance the efficiency of consumers’ green product purchase through a cognitive process that involves early attentional congruency. Practically, the present research provides valuable insights for retailers aiming to increase sales of green products.

## 2. Theoretical backgrounds and hypotheses

The S-O-R model, derived from environmental psychology by Mehrabian and Russell in 1974 and further conceptualized by Jacoby in 2002, provides a broader perspective compared to solely focusing on marketing effectiveness. This approach considers both external and internal factors that influence consumer responses (Huang et al., 2021; Marikyan & Papagiannidis, 2023). Specifically, the S-O-R model conceptualizes external factors, such as ambient store sounds (Zheng et al., 2019). It considers the internal psychological or physiological processes of consumers as the organismic component (Jiang et al., 2020), and consumer attitudes and behaviors towards products are regarded as responses within this model (Chen et al., 2023; Bradford & Desrochers, 2009). Next, we focused on “stimulator”, “organism” and “response” in this study.

### 2.1. Natural sounds and purchase decision of green products

A direct response to the stimulus of natural sounds promoting green consumption is an increase in purchase rate for green products. Previous research has indicated that natural sounds have the potential to enhance male consumers’ willingness to purchase climate-friendly and organic foods (Spendrup et al., 2016). Additionally, natural sounds have been found to increase consumers’ choice of healthier food options (Peng-Li et al., 2021; 2022). Furthermore, Ferreira and Oliveira-Castro (2011) observed a significant difference in actual expenditures between conditions with classical music, which resulted in higher spending compared to both the no music and pop music conditions. Similarly, in a study conducted by North and Hargreaves (1998), it was found that both classical and pop music increased actual cafeteria sales when compared to easy listening and quiet music. Therefore, based on the aforementioned findings, we can infer the following hypothesis:

**Hypothesis 1a** Natural sounds (vs. unnatural sounds) have a significant positive impact on the purchase rate for green products (vs. non-green products).

The response of consumers to natural sounds associated with green products can also be reflected in the efficiency of the evidence accumulation process in purchase decision-making (Evans, 2008; Krajbich et al., 2012). This efficiency not only includes purchasing more green products but also involves purchasing them more quickly. Peng-Li et al. (2022) did not find a significant difference in the response time between the choice of healthy and unhealthy food options under natural sounds. This could be attributed to their exclusive focus on response time, while overlooking choice frequency, which may not accurately reflect whether consumers are selecting a particular product more quickly. To comprehend the efficiency of the evidence accumulation process, we opted for the HDDM.

The HDDM is a model that captures decisions rooted in value-based decision theory (Wiecki et al., 2013; Ratcliff et al., 2016). HDDM, based on Bayesian framework, quantifies uncertainty using observed data and a prior distribution, rather than relying solely on a theoretical distribution as in classical frequentist analyses. Moreover, within the Bayesian framework, this model enables the measurement of decision process speed by taking into account factors such as accuracy (Vandekerckhove et al., 2011). Hence, the HDDM allows for the joint modeling of both response time and choice patterns (purchase or no-purchase) across trials to infer the efficiency of evidence accumulation. This marks an advancement over simpler models that solely concentrate on modeling response time or choice patterns.

As illustrated in Fig. 1, it is apparent that the HDDM classifies “purchase” and “no-purchase” as upper and lower bound responses, respectively. As time progresses, a drift process occurs. During this drift, consumers begin to accumulate evidence, eventually causing their choices to gradually converge towards a certain boundary. Thus, the consumer reaches the decision (Ratcliff & Rouder, 1998; Smith & Ratcliff, 2004; Wiecki et al., 2013). A higher drift rate implies more efficient accumulation of evidence for purchasing decisions, as well as a higher probability of a shorter reaction time when choosing to purchase (vs. not purchasing).

**Fig. 1.**
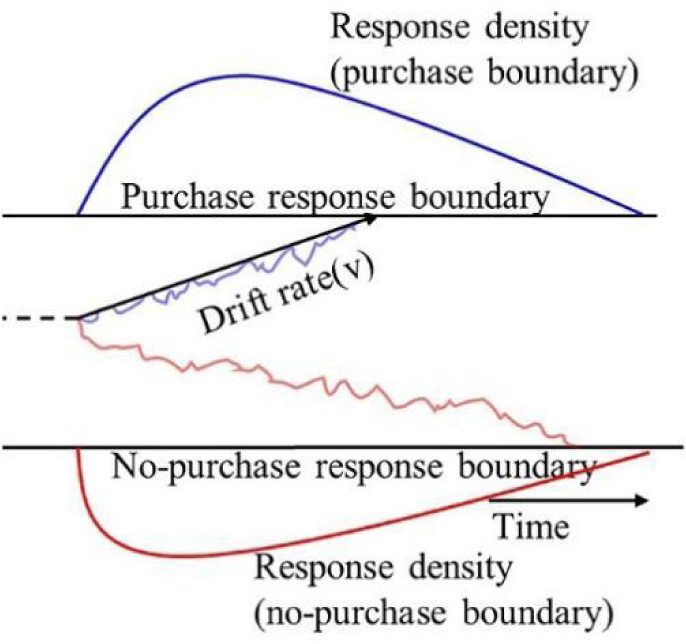
The visualization of the drift rate for HDDM. The blue line within the boundary symbolizes the evidence accumulation process for purchasing, while the red line represents the same process for not purchasing. The panels labeled “purchase” (blue) and “no-purchase” (red) encompass density plots at the boundaries of the two possible responses over time.

Previous studies have demonstrated that background sounds can impact the speed of consumption (Milliman, 1982; Oakes, 2000). For instance, background sounds with a fast tempo have been shown to expedite consumers’ decision-making in a store (Smith & Curnow, 1966) and can also hasten the pace at which consumers eat (Caldwell & Hibbert, 2002). Therefore, our hypothesis is that natural sounds will increase the drift rate in consumer purchase decisions for green products. The specific hypotheses are as follows:

**Hypothesis 1b** Natural sounds (vs. unnatural sounds) significantly enhance the efficiency of purchase decisions (drift rate) for green products (vs. non-green products).

### 2.2. Natural sounds and early attentional congruency

Now, let’s shift our focus to the “organism” component within the S-O-R model. The impact of natural sounds on the processing of information related to green products may be reflected in the early attentional mechanisms. Drawing upon the attention restoration theory, it has been observed that natural stimuli reduce the demand for attention (Kaplan, 1995). Furthermore, based on an evolutionary perspective, there exists an instinctual connection between humans and nature (Kellert & Wilson, 1993). Natural sounds may have quicker and subtler effects on consumers’ processing of information about green products, which could be reflected in early attention mechanisms involved in product information processing. ERP techniques provide a unique opportunity to uncover early attentional mechanisms involved in the processing of product information, given their excellent temporal resolution (Yun et al., 2022). For instance, Ozkara and Bagozzi (2021) found that early ERP components (before 300 milliseconds [ms]) can reflect consumers’ automatic and rapid cognitive processes during the processing of product information.

This early attention mechanism may be reflected in whether consumers perceive the congruency between the sound environment and the product. In actual marketing, dealers often use background sounds to create a specific retail atmosphere (Anglada-Tort et al., 2022; Ferreira & Oliveira-Castro, 2011). According to the musical hypothesis, consumers are activated by background sounds that are congruent with relevant information, leading them to choose products that are congruent with the background sounds (Alpert & Alpert, 1990; North et al., 2016). A large body of research has also demonstrated that when background sounds and a product are congruent, they enhance the consumer’s choice of that product (Helmefalk & Hultén, 2017; Jacob, 2006; Morrin & Chebat, 2005). Therefore, natural sounds may influence consumers’ early attentional congruency when processing information about green products.

Early attentional congruence is reflected in the P2 component, which is a positive potential with a latency of 200 ms. The P2 component typically peaks at the centro-parietal regions of the brain (Dong et al., 2010; van Hooff et al., 2011). In previous studies, the P2 component has been found to be significantly reduced in congruent stimuli (Freunberger et al., 2007; Gruber & Müller, 2005), including sound stimuli (Zioga et al., 2020; Debnath & Wetzel, 2022). The P2 component was also found to be lower in response to high-order online shopping webpages compared to low-order online shopping webpages (Shang et al., 2020). This may be due to the lower demand for integrating visual information and coping with the sense of congruency in the high-order layout. Natural sounds can also strengthen a consumer’s connection with nature (Buxton et al., 2021), which may increase consumers’ perception of the congruency between green products and the environment. Therefore, we hypothesize that the P2 component, as a reflection of early attentional congruence, is associated with the influence of natural sounds on the processing of information related to green products.

In summary, we hypothesize that:

**Hypothesis 2** Natural sounds significantly influence early attentional congruency during the processing of information regarding green products. More specifically, natural sounds (vs. unnatural sounds) significantly reduce the P2 component during the processing of information regarding green products (vs. non-green products).

### 2.3. Eearl attentional congruency related to decision-making efficiency

Attentional mechanisms are crucial in enhancing the efficiency of purchase decision-making, as demonstrated by corresponding neural signals. As previously mentioned, neural signals can elucidate the mechanisms of consumer attention when processing product information (Ozkara & Bagozzi, 2021; Shang et al., 2020). Furthermore, neural signals can also serve as predictors of purchase decisions (Ozkara & Bagozzi, 2021; Telpaz et al., 2015). In a previous study, researchers found that electroencephalogram (EEG) signals serve as indicators of the evidence accumulation process (Pisauro et al., 2017). Futhermore, Liu and Wang et al. (2022) unveiled a correlation between the ERP components associated with attentional mechanisms and the efficiency of evidence accumulation during decision-making. Additionally, Mennella et al. (2020) discovered that the efficiency of evidence accumulation is linked to early attentional mechanisms, as manifested in EEG components occurring between 145 and 265 ms.

Decision efficiency is not only associated with early attention, but also related to congruency. A study examining the congruency between background sounds and products in terms of social identity found that consumers exhibited a higher willingness to pay within a short timeframe (North et al., 2016). Based on this evidence, it can be inferred that greater congruence in perception influences the speed of consumer decision-making.

Taken together, we propose the following hypothesis:

**Hypothesis 3** The influence of natural sounds on the decision efficiency of green products is mediated by early attentional congruency during product information processing. Specifically, the presence of natural sounds (vs. unnatural sounds) enhances the drift rate towards selecting green products (vs. non-green products) by reducing the P2 component during the processing of product information.

In conclusion, our study seeks to examine the impact of natural sounds on influencing consumers’ inclination to purchase green products. We adopt the S-O-R model as our theoretical framework to explain how natural sounds nudge the efficiency of decision-making related to green product purchase. This enhancement is attributed to the facilitation of early attentional congruency by natural sounds during the processing of information pertaining to green products (see Fig. 2).

**Fig. 2.**
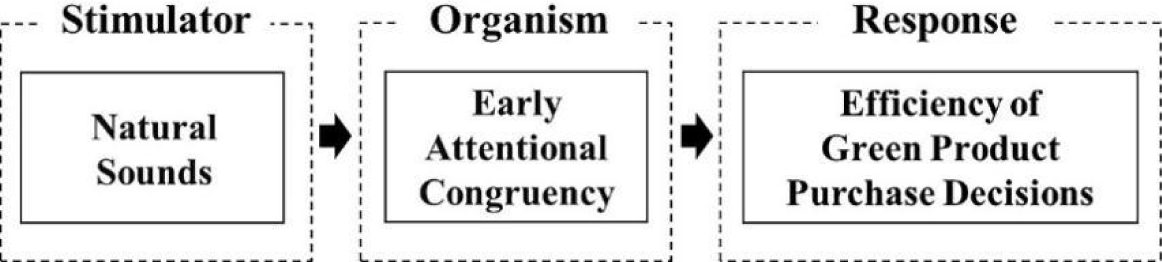
The research conceptual model.

## 3. Method

### 3.1. Participants and design

The G*Power software analysis suggested that we need to collect at least 24 participants to detect a reliable effect (*α* = 0.05, 1-*β* = 0.80, ANOVA: Repeated measure, within factors, Faul et al., 2009). Conservatively, 26 participants aged from 19 to 27 years (16 males, with a mean age [M] of 21.65 years and a standard deviation [SD] of 1.90 years) were recruited. All participants reported having no history of psychiatric illness, normal or corrected-to-normal vision, and being right-handed. Additionally, all participants ensured they had enough rest and had not taken any medications recently. Participants gave written consent to participate in this experiment, and they also received the payment or products due to them for their participation. The study was approved by the local ethics committee and was in line with the Helsinki Declaration of 1964.

This experiment used a 2×2 within-subject experimental design, with factors including sound conditions (natural vs. unnatural) and product types (green vs. non-green), to test our hypothesis.

### 3.2. Materials

#### 3.2.1. Products

The experiment sampled 54 products from https://www.jd.com/, which is one of China’s largest e-commerce sites. Each product included two versions of the green China green product (CGP)^1^label indicating that the products were green and the grey CGP label indicating that the products were non-green (Linder et al., 2010; Schwartz et al., 2020). Refer to Griskevicius et al. (2010), Green products were described as superior in terms of pro-environmental benefits, while non-green products were described as superior in terms of performance (see Appendix for details). Also, referring to Griskevicius et al. (2010), the two types of products were always priced equally. Additionally, referring to Knutson et al. (2007), we applied a 75% discount to the website price of these products to encourage purchases, resulting in participants seeing prices ranging from 2.25 RMB to 16.5 RMB during the experiment. Referring to Knutson et al. (2007), out of these 54 products, 27 were in the inexpensive category (less than 10 RMB), while the other 27 were in the expensive category (more than 10 RMB) (see Table A1 for details). To ensure that these products were everyday necessities, the likelihood of purchase and familiarity for all products was higher than the median (see Table A1 for details).

#### 3.2.2. Sounds

The sound of birds chirping was selected as the representative for natural sounds (Spendrup et al., 2016), while machine noise was chosen to represent unnatural sounds (Koivisto et al., 2022). All sounds were selected from a free sound library available at https://freesound.org/. Each sound segment’s length coincided with the maximum duration of a single trial, approximately 10.6 seconds, and was played repeatedly during the experiment. The sound intensity was uniformly set to one level using Adobe Audition software and was tested to confirm a playback volume of 50 dB using a decibel detector. The processed sound files can be accessed at https://osf.io/3754a/. The processed sounds were rated on a 9-point scale for natural and unnatural sounds (1=more relaxing; 9=more arousing) by an additional 10 participants (*Mage* ± SD = 25.7 ± 1.35, 6 males) who did not participate in the formal experiment (Peng-Li et al., 2022). The results indicated that participants experienced greater relaxation in response to natural sounds (3.60 ± 1.26) compared to unnatural sounds (6.60 ± 0.84), *t* (9) = −7.115, *p* < 0.001, Cohen’s *d* = 2.259.

### 3.3. Experiment procedure

This experimental task was adapted from the SHOP (i.e., “Save Holdings Or Purchase”) task originally developed by Knutson et al. (2007). As depicted in Fig. 3, following the cross-point gaze, participants proceeded to the initial stage of product information (Stage 1), where a picture and name of the product would be displayed. Participants were instructed to simply view the product image for a duration of 2 seconds, without providing any response. After progressing to the second stage of product information (Stage 2), participants were not required to provide any response even when viewing the price for a duration of 2 seconds. Upon reaching the subsequent screen, participants entered the decision-making period, during which a screen displaying the options “buy” and “not” was presented. Half of the participants were instructed to press the “F” key to indicate “buy” and the “J” key to indicate “not”. In the other half of the participants, they were instructed to press the “J” key to indicate “buy” and the “F” key to indicate “not”. If the participant did not respond within 2 seconds, a new identical trial was repeated until a suitable response was made. Additionally, if participants pressed the key faster than 200ms, it was considered a response that was too quick and they were required to repeat that trial. To incentivize participants to participate in the SHOP task, they were initially given an explanation of the experiment. They were informed that we would randomly select one of their choices and provide a cash or product reward for it. In the event of compliant choice button responses, selecting “buy” would result in the cashing out of the remaining funds and the acquisition of the product. Conversely, selecting “not” would lead to the cashing out of the principal. However, if a participant failed to respond within 2 seconds or responded faster than 200 milliseconds during the choice phase, the session would be voided and they would forfeit their 20 RMB principal and the opportunity to purchase the product. Prior to the commencement of the formal experiment, participants completed four practice trials without any record of their choices. Upon confirming their understanding of the rules, they proceeded to the formal experiment.

**Fig. 3.**
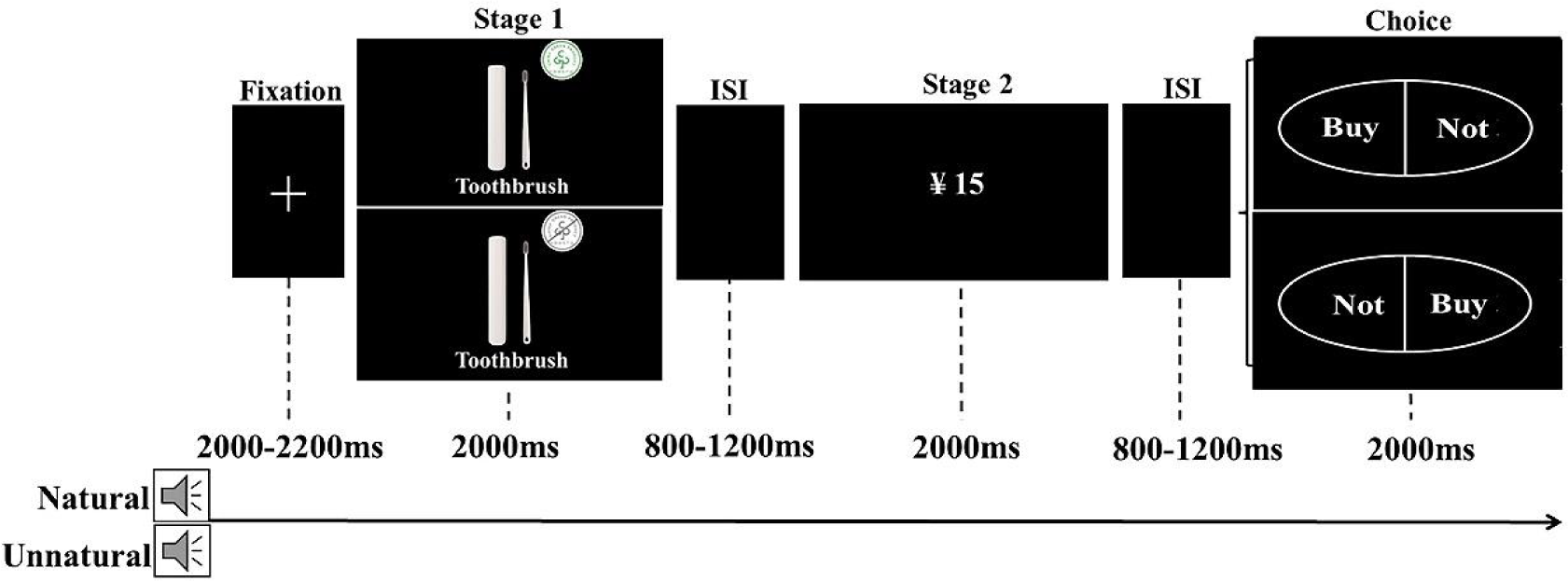
Time sequence of stimuli in each trial of the SHOP task. ISI= interstimulus interval. The products were divided into two sets of 54 products, with each set containing 27 products.

These stimulus sets were counterbalanced, meaning that both sets were presented an equal number of times in the presence of natural and unnatural sounds between participants. Participants were randomized into one of two orders, either hearing natural sounds first or unnatural sounds first. Participants completed a total of 6 times, with three times conducted under the unnatural sound condition and three times under the natural sound condition. Each time included 27 green products and 27 non-green products. The order of green and non-green products was balanced within each block. In each trial, the order of product was randomized to prevent participants from forming expectations (Zhang et al., 2021). At the conclusion of each sound condition, participants provided ratings for the sound in that condition as a manipulation check, using a 9-point scale (1=more relaxing; 9=more arousing; Peng-Li et al., 2022). After completing all experimental tasks, participants provided ratings on six items to assess product greenness and product practicability as manipulation checks (product greenness: *α*green label = 0.88, *α*non-green label = 0.87; product practicability: *α*green label = 0.87, *α*non-green label = 0.95, see Appendix, adapted from Yan et al., 2021).

### 3.4. Experiment apparatus and data analysis

#### 3.4.1. Electroencephalogram recording and analysis

In this experiment, 32 Ag/AgCl scalp electrodes positioned at the standard 10–20 locations were used to collect raw EEG data (Neuroscan, Neurosoft Labs, Inc. Sterling, USA). Additionally, vertical electrooculogram (EOG) activity was recorded with electrodes placed supra- and infra-orbitally at the left eye. Horizontal EOG activity was recorded with electrodes placed on the left and right orbital rim (Wang et al., 2021). All leads were referenced to the bilateral mastoids, and the electrode impedance was kept at about 10 kΩ.

The collected data was analyzed in MATLAB using the EEGLAB toolkit. The signals were filtered using Butter-worth filters with half-power cutoffs at 0.01 and 30 Hz and down-sampled at a rate of 500 Hz (Zhang et al., 2021). All trials were de-artifacted within the range of ±100 μV. Finally, there were 26 participants with an average artifact rejection rate of 11.17%. Each of our analytical conditions can satisfy the trial requirements (Luck, 2005; see Appendix descriptions for details). The timeline for ERP analysis was set from 200ms before to 800ms after the onset of stimuli, applied to both Stage 1 and Stage 2 of the product information progression, respectively. Based on the visual inspection and prior research (Goto et al., 2017), we extracted mean amplitudes of the time windows: 150–230 ms (P2). We selected eleven electrodes on the forehead (FP1, FP2, F3, FZ, F4 FC3, FCZ, FC4, C3 CZ and C4) to define a specific area of interest (Goto et al., 2017; Lee et al., 2014).

#### 3.4.2. Data analysis

We conducted HDDM analysis of experimental behavioral data from 26 participants (Wiecki et al., 2013). The HDDM analysis was conducted on the Python platform (Wiecki et al., 2013). Following the fitting of the model, we utilized Monte Carlo Markov Chain simulation (MCMC) with gradient ascent optimization to approximate the posterior distributions of the model parameters. We took 10,000 samples and burned the first 1,000 to stabilize the model (Regenbogen et al., 2016). We ran the simulation five times for each model in order to assess convergence (Wiecki et al., 2013). In addition, we used Gelman-Rubin (R^) as an evaluation value for convergence (Gelman & Rubin, 1992). The closer R^ values are to 1 indicates better convergence.

We used Statistical Package for the Social Sciences (SPSS) 23 to process behavioral and ERP data through conducting the paired-samples t-tests, repeated measures analysis of variance (ANOVA) and mediation analysis.

## 4. Result

### 4.1. Manipulation checks

A paired-samples t-test on participants’ reported feelings about the sound conditions revealed that the natural sound was perceived as significantly more relaxing than the unnatural sound, *M* natural ± SD = 3.15 ± 1.29 vs. *M* unnatural ± SD = 6.81 ± 1.17, *t* (25) = −10.838, *p* < 0.001, Cohen’s *d* = 2.122. A paired-samples T-test on the ratings for product greenness revealed that products with a green label were significantly rated higher than products with a non-green label in terms of product greenness *M* green label ± SD = 5.60 ± 1.06 vs. *M* non-green label ± SD= 2.19 ± 0.81, *t* (25) = 10.812, *p* < 0.001, Cohen’s *d* = 2.127. A paired-samples T-test on the ratings for product practicability revealed that products with a green label were significantly rated lower than products with a gray label in terms of product practicability, *M* green label ± SD = 3.32 ± 1.11 vs. *M* non-green label ± SD = 4.91 ± 1.53, *t* (25) = −4.122, *p* < 0.001, Cohen’s *d* = 0.807. Therefore, the manipulations of sound conditions and product types were both successful.

### 4.2. Behavioral date

#### 4.2.1. Purchase rate

To test the hypothesis that the natural sound increases the purchase rate for green products, a 2×2 (Sound conditions) × (Product types) repeated measure ANOVA^2^ was conducted, with the purchase rate as the dependent variable and order of the sounds (natural sound first/unnatural sound first) as the covariate. The interaction effect was significant, *F* (1,24) = 5.838, *p* = 0.024, 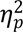 = 0.196. Pairwise comparison^3^ indicated that, as illustrated in Table 2 and Fig. 4, there was a significant effect of sound conditions on the purchase rate for green products, but an insignificant effect for non-green products. The main effect of product type was significant, *F* (1,24) = 5.147, *p* = 0.033, 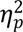 =0.177, *M* green products ± SD = 0.38 ± 0.17, vs. *M* non-green products ± SD = 0.33 ± 0.16. The other effects did not reach significance at the 0.1 level.

**Fig. 4.**
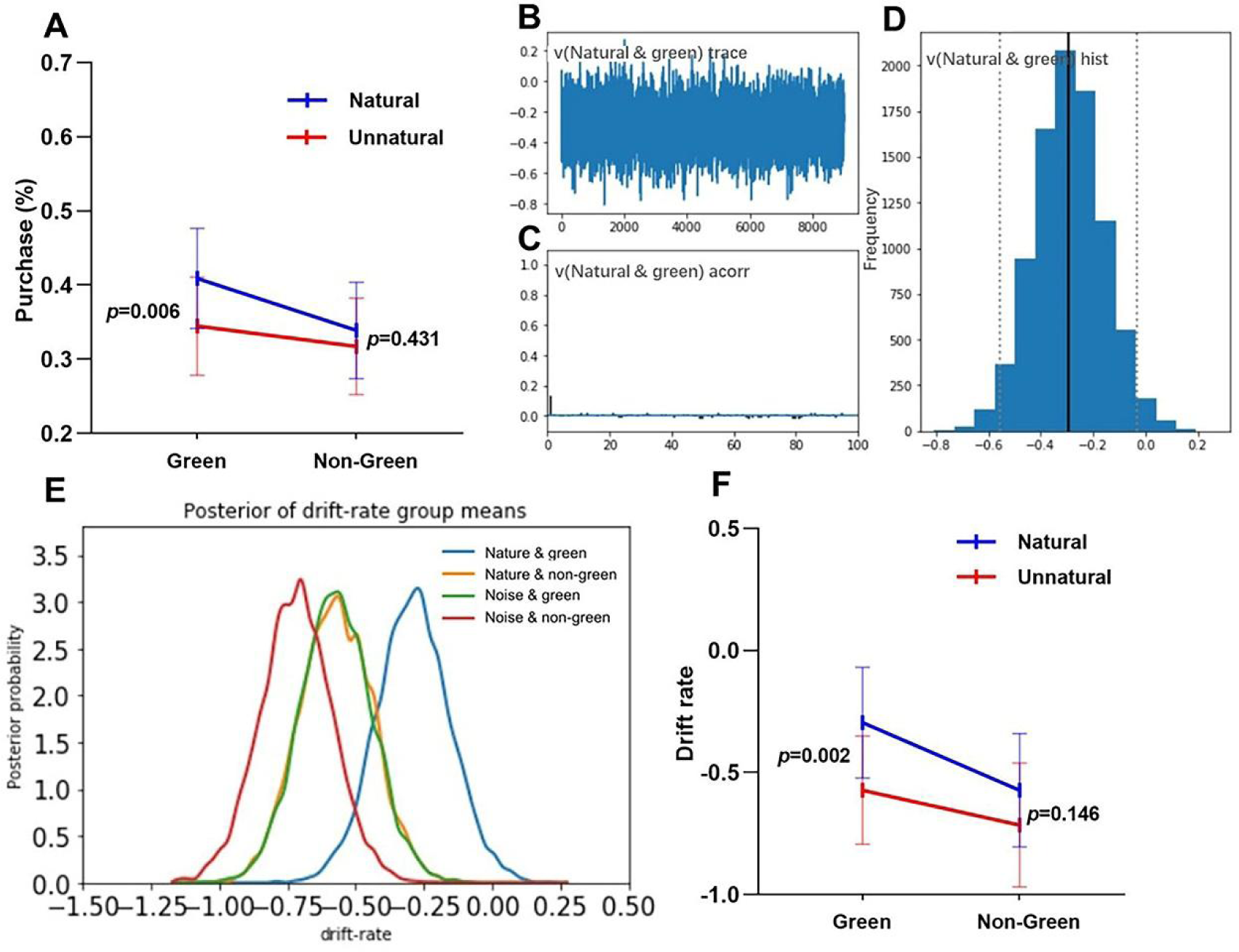
Results for behavioral data. (A) Line chart of the mean and 95% confidence interval (CI) of the purchase rate across the 2 (Sound conditions) by 2 (Product types) conditions. (B)The trace plot for 9000 iterations. (C) The autocorrelation for the last 100 iterations. (D) The histogram of the drift rate value. (E) Posterior probability densities for the 2 (Sound conditions) by 2 (Product types) conditions estimated from the HDDM by drift rates. Peaks reflect the best estimates, while width represent uncertainty. (F) Line chart of the mean and 95%CI of the drift rates across the 2 (Sound conditions) by 2 (Product types) conditions.

**Tabel 2.** Interaction effect on purchase rate and drift rate.

| Behavioral metrics | Product types | Sound conditions | M±SD | $F(1,26)$ | $\eta_p^2$ |
| --- | --- | --- | --- | --- | --- |
| Purchase rate | Green | Natural | 0.41±0.17 | 8.888** | 0.270 |
| | | Unnatural | 0.34±0.16 | | |
| | Non-green | Natural | 0.34±0.16 | 0.643 | 0.026 |
| | | Unnatural | 0.32±0.16 | | |
| Drift rate | Green | Natural | -0.30±0.56 | 12.475** | 0.342 |
| | | Unnatural | -0.57±0.55 | | |
| | Non-green | Natural | -0.57±0.58 | 2.254 | 0.086 |
| | | Unnatural | -0.72±0.62 | | |

<sup>†</sup> $p < .1$ ; \* $p < .05$ , \*\* $p < .01$ , \*\*\* $p < .001$ .

#### 4.2.2. HDDM analysis

##### 4.2.2.1. Model convergence

As depicted in Fig. 4, considering one condition of the drift rate as an example (others see Fig. A1 in the Appendix), we ascertained that the model exhibits a cohesive trajectory characterized by smoothness. Furthermore, the autocorrelation tended towards zero, indicating minimal dependency between subsequent data points, and the distribution of values was normal. The Gelman-Rubin statistics (R^) for the model were very close to 1 (mean = 1.00002; SD = 0.00006), indicating that the convergence performance of our model was satisfactory.

##### 4.2.2.2. Drift rate

To test the hypothesis of natural sounds on the efficiency of green product purchase, we conducted a 2×2 (Sound conditions) × (Product types) repeated measures ANOVA with drift rate as the dependent variable and the order of the sounds as the covariate.

The interaction effect was significant, *F* (1,24) = 4.993, *p*= 0.035, 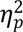 = 0.172. Pairwise comparison indicated that, as illustrated in Table 2 and Fig. 4, there was a significant effect of sound conditions on the drift rate for green products, but an insignificant effect for non-green products. The main effect of product type was also significant, *F* (1,24) = 7.092, *p* = 0.014, 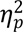 = 0.228. Then drift rate for green products was significantly higher than non-green products, *M* green products ± SD =-0.43±0.57 vs. *M* non-green products ± SD=-0.64±0.60. The other effects did not reach significance at the 0.1 level.

### 4.3. ERP date

To test the hypothesis that natural sounds influence early attentional congruency during the processing of green product information, we conducted 2×2 (Sound conditions) × (Product types) repeated measure ANOVA with P2 components as the dependent variable and order of the sounds as the covariate.

In Stage 1, the main effect of sound conditions was significant, *F* (1,24) = 7.763, *p*=0.010, 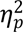 = 0.244, *M* natural ± SD = −0.21 ± 2.29 vs. *M* unnatural ± SD = 0.40 ± 2.02. The other effects did not reach significance at the 0.1 level.

In Stage 2, the results showed a marginally significant interaction between sound conditions and product types, *F* (1,24) = 3.791, *p* = 0.063, 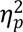 = 0.136. Decomposing the interaction, as illustrated in Table 3 and Fig. 5, we observed a significant effect of sound conditions on the mean amplitude of P2 for green products, but an insignificant effect of sound conditions for non-green products. The main effect of sound conditions showed marginal significance, *F* (1,24) =3.303, *p*=0.082, 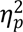 = 0.121, *M* natural ± SD =0.83 ±2.62 vs. *M* unnatural ± SD = 1.87 ±2.25. The other effects did not reach significance at the 0.1 level.

**Fig. 5.**
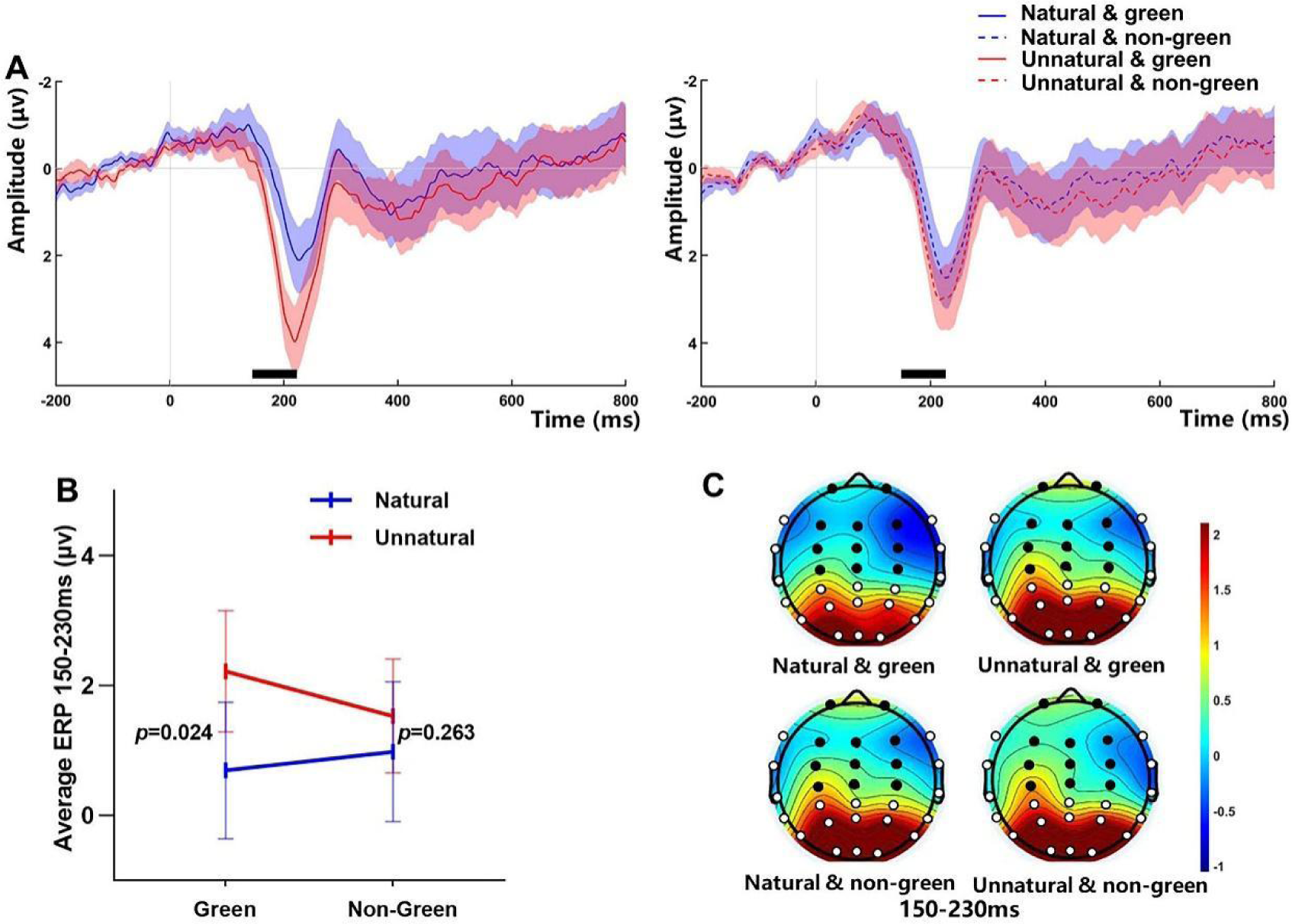
The interaction effect of sound conditions and product types on mean amplitude of the P2 (Stage 2). (A) The average ERP waveforms during the 2 (Sound conditions) by 2 (Product types) conditions for channels (FP1, FP2, F3, FZ, F4 FC3, FCZ, FC4, C3 CZ and C4). Intensity-dependent differences are identified during an early (i.e., 150–230 ms) window (marked with a blank bar). Shaded areas represent the standard error of the mean for the ERP signal at each time point. (B) Line chart of the mean and 95%CI of the P2 amplitude across the 2 (Sound conditions) by 2 (Product types) conditions. (C) Scalp topography during the early window after the vibration onset relative to the pre-stimulus baseline for the 2 (Sound conditions) by 2 (Product types) conditions. The eleven dots marked in black are FP1, FP2, F3, FZ, F4 FC3, FCZ, FC4, C3 CZ and C4 channels.

**Table 3.** Interaction effect of sound conditions and product types.

| Product information | Product types | Sound conditions | $M \pm \text{SD}$ | $F(1,24)$ | $\eta_p^2$ |
| --- | --- | --- | --- | --- | --- |
| Stage 2 | Green | Natural | $0.69 \pm 2.60$ | 5.767* | 0.194 |
| | | Unnatural | $2.22 \pm 2.32$ | | |
| | Non-green | Natural | $0.98 \pm 2.67$ | 1.315 | 0.052 |
| | | Unnatural | $1.53 \pm 2.17$ | | |

### 4.4. Mediation analysis

To test the hypothesis that natural sounds, as opposed to unnatural sounds, impact early attentional incongruency, which in turn affects the drift rate for green products, we conducted a mediation analysis. This analysis utilized 5,000 bootstrap samples (Model 4, Hayes, 2013). The independent variable was sound conditions, the mediator was the P2 of Stage 2, the dependent variable was the drift rate for green products, and the covariates included the order of sounds, age, and gender. As illustrated in Fig. 6, the findings indicated that the P2 of Stage 2 could account for the promotional impact of sound conditions on drift rate for green products, *b*=0.114, *SE*=0.072, 95%CI = [0.001, 0.275]. Additionally, we performed a similar mediation analysis, with the drift rate for non-green products as the dependent variable. The indirect effect did not exclude zero, *b*=0.014, *SE*=0.036, 95%CI = [-0.029, 0.112].

**Fig. 6.**
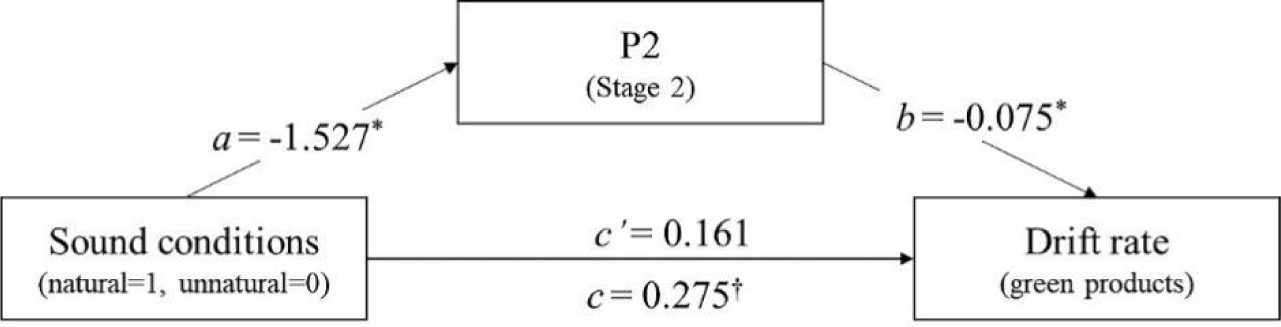
Mediation analysis for sound conditions on drift rate.

## 5. Discussion

### 5.1. Findings

The present study provides new insights into how natural sounds can nudge green consumption, building upon the S-O-R model (Mehrabian & Russell, 1974; Jacoby, 2002). Specifically, the current study employed the HDDM analysis and ERP technology to discover that natural sounds increased early attentional congruency in processing green product information, thereby leading to more and faster decisions to purchase green products.

Firstly, consistent with the findings of the previous studies conducted by Spendrup et al. (2016) and Peng-Li et al. (2021; 2022), it is evident that natural sounds have the capacity to nudge green consumption. This finding provides support for our hypothesis 1a. To further explore the mechanism by which natural sounds impact purchasing decisions for green products, the current study utilized the HDDM analysis method. The findings revealed that natural sounds had a significant impact on the drift rate linked to decisions to purchase green products. This finding provides support for our hypothesis 1b. This result suggests that natural sounds may impact the efficiency of consumers’ decision-making process when purchasing green products. Differing from the methodologies employed in extant literature (Smith & Curnow, 1966; North et al., 2016), this finding introduces an innovative application of drift rate as a metric to validate previous findings that demonstrate the influence of sound on the speed of consumer decision-making.

Secondly, during the processing of product information, we observed in the first stage that consumers demonstrated higher levels of early attentional congruency when exposed to natural sounds. Furthermore, in the second stage, we found that consumers exhibited higher levels of early attentional congruency towards green products when exposed to natural sounds. These results partly support our hypotheses 2. Differing to the visual attention mentioned previously (Peng-Li et al., 2021), the current study leverages the temporal advantages of ERP technology to uncover an enhancement in early attentional congruency during the processing of information related to green products induced by natural sounds. The observed increase in early attentional congruency in brain activity during the processing of green products, influenced by natural sounds, can potentially be attributed to several factors. These include the regulation of attentional resources by natural cues (Kaplan, 1995), consumers’ evolutionary preference for bird sounds as mentioned in an evolutionary perspective (Kellert & Wilson, 1993), and the impact of congruency between background sounds and the products (Alpert & Alpert, 1990; North et al., 2016). In addition, these results suggest that participants would show a lower P2 wave towards green products only after all product information (including Stages 1 and Stages 2) has been presented. This could be attributed to participants receiving information that their purchasing decisions would be randomly redeemed, which in turn led them to exercise greater caution.

Thirdly, we observed that an increase in early attentional congruency during Stage 2 of the processing of product information, facilitated by natural sounds, was associated with an increase in drift rate for green products. This result provides partly support for our hypothesis 3. This finding is consistent with previous research linking attention-related brainwave components and decision-making efficiency (Liu and Wang et al., 2022; Mennella et al., 2020) as well as the impact of perceptual congruency on consumer decision making (North et al., 2016). It suggests that attention mechanisms may be a critical factor influencing decision-making efficiency. Moreover, what sets this study apart from previous research is its demonstration of the impact of natural sounds on early attentional congruency in the context of shopping, thereby improving the efficiency of purchase decisions for green products. This finding further enriches our understanding of how natural sounds can nudge individuals towards green consumption.

### 5.2. Theoretical and practical implications

Based on the S-O-R model, this study introduces an early attention congruency to elucidate the nudge effects of natural sounds on the decision efficiency of evidence accumulation process in green consumption. This contribution offers significant theoretical and practical implications.

In terms of theoretical implications, firstly, the majority of research on the S-O-R model has focused on exploring indicators of willingness to respond (Fei et al., 2021; Liu and Liu et al., 2022; Tandon et al., 2023). The findings of this study indicate that natural sounds not only boost the purchase rate of green products but also enhance the efficiency of the decision-making process for purchasing green products. This approach expands the scope of response indicators based on the perspective of evidence accumulation. Secondly, in this study, we found that natural sounds facilitated early attentional congruency in consumers’ processing of information for green products. This enriches the existing literature on the influence of natural sounds on attention during product information processing from a temporal dimension (Peng-Li et al., 2021). Thirdly, within the framework of the S-O-R model, this study identified a link between attention-related ERP components under the influence of natural sounds and the drift rate. This innovative approach provides new perspectives on the application of the S-O-R model for investigating neural processes and behavioral responses. It also offers empirical evidence for mapping physiological and psychological concepts to computational parameters (Roberts & Hutcherson, 2019).

In terms of practical implications, firstly, this study chose to focus on incentive purchase decisions (Knutson et al., 2007), making it a more realistic and applicable choice. This suggests that natural sounds can indeed be effectively utilized in real shopping scenarios to encourage consumers to choose green-labeled products. Secondly, in previous studies focusing on marketing effectiveness, few scholars have paid attention to the indicator of decision efficiency. Based on the perspective of natural sounds promoting green consumption, this study has raised awareness of decision efficiency for green products. This, to a certain extent, provides a richer set of marketing effectiveness assessment indicators to evaluate marketing effectiveness in practical markets. It also offers more in-depth evidence to guide marketers in using natural sounds to promote green consumption. Thirdly, the results of this study suggest that marketers should also focus on consumer brain activity and product information when formulating marketing strategies (Stanton et al., 2017; Robertson et al., 2017). When consumers perceive a harmonious relationship between green products and the surrounding environment in their brains, it further promotes their green consumption. Based on this guiding principle, marketers can not only use natural sounds to create harmony between the environment and green products, but also utilize natural landscapes and other measures to better promote green consumption.

### 5.3. Limitation and future research outlooks

The current study has the following limitations. Firstly, it is worth mentioning that this study specifically utilized bird sounds as the natural sound stimulus. However, it is important to note that there are numerous other natural sounds found in nature, such as the sounds of waves (Peng-Li et al., 2022), waterfalls (Koivisto et al., 2022), and rain (Lin et al., 2022). Future research could further investigate the potential of using these sounds in marketing strategies to promote green consumption. Secondly, this study specifically focused on analyzing ERP, which are indicators of neural activity in the brain. The ERP was examined to uncover the mechanisms through which natural sounds enhance the effectiveness of purchasing decisions for green products. In future studies, other physiological metrics such as heart rate and skin conductance (Xu et al., 2019) could be included to enhance the comprehensive understanding of the effects of natural sounds on consumer behavior. This approach would lead to a deeper understanding of the underlying mechanisms involved. Thirdly, it is important to note that this study solely analyzed data from consumers performing tasks within a laboratory setting. However, it is important to consider that the temporal effects describe the persistence of the response over time (Reimann et al., 2012). Prior research has shown that connecting with nature has a long-lasting impact on thoughts and behaviors, including fostering pro-environmentalism (Chen et al., 2023), and promoting green consumption (Cooper et al., 2015). Therefore, future research could consider conducting a longitudinal investigation to examine the sustained promotional effect of natural sounds on green consumption.

## Appendix

### Specific descriptions for products

Green products are characterized by their sustainable access to resources, environmentally friendly production, marketing strategies, usage, and disposal throughout all stages of their life cycle. They are notable for their low resource consumption, minimal pollutant emissions, and ease of recycling and reuse. In contrast, non-green products are characterized by higher standards in terms of materials, structure, and functionality. These products prioritize practicality, efficiency, and reliability.

**Table A1.**
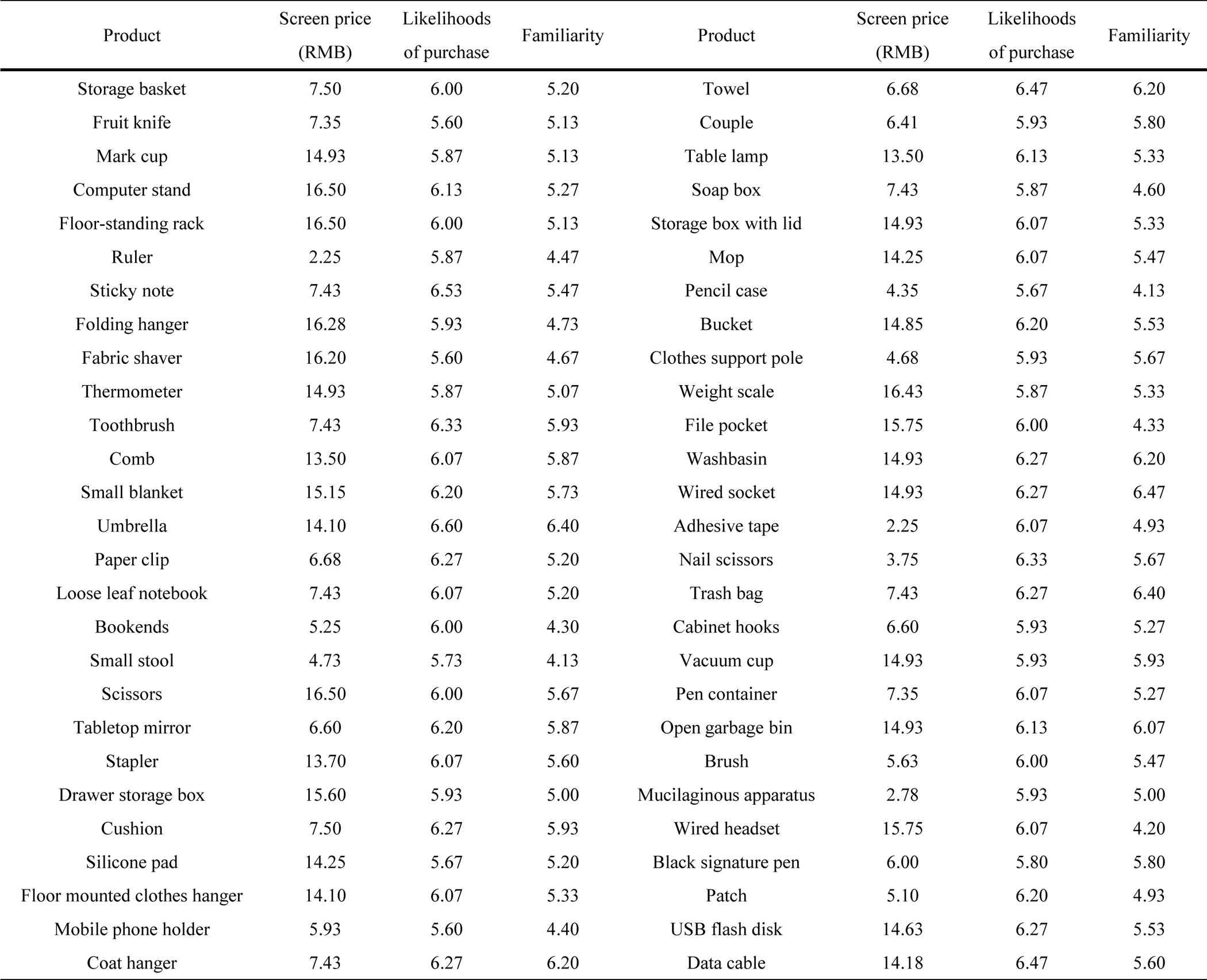
Product name, screen price, likelihoods of purchase and familiarity.

### Likelihoods of purchase and familiarity ratings

Purchase likelihood and familiarity were assessed on a 7-point scale (1= not at all, 7= extremely). The first assessment was rated by 15 participants (*Mage* ± SD = 25.33 ± 1.92, 10 males) who were not involved in the formal experiment. After the first assessment, 47 products that scored above the median in both likelihood of purchase and familiarity were selected. The second assessment was rated by 15 other participants (*Mage* ± SD = 25.33 ± 2.17, 9 males) who were not involved in the formal experiment. After the second assessment, 7 products that scored above the median in both likelihood of purchase and familiarity were selected.

### Product greenness and product practicability

Participants were asked to what extent they agreed with the following statements under green products label and non-green products label (1=strongly disagree, 7= strongly agree):

Products with this label are environmentally friendly.
Products with this label are green.
Products with this label are good for the environment.
Products with this label are practical.
Products with this label are efficient.
Products with this label are reliable.

### Specific number of trials to analyze conditions

During the first stage of product information processing, each participant viewed green products in

71.00 ± 15.54 trials and non-green products in 69.60 ± 16.32 trials under natural sound conditions. Under unnatural sound conditions, there were 72.88 ± 12.80 trials for viewing green products and 71.68 ± 14.00 trials for viewing non-green products per participant. During the second stage of processing, participants viewed the prices of green products in 71.52 ± 14.33 trials and the prices of non-green products in 71.52 ± 14.33 trials under natural sound conditions. Under unnatural sound conditions, they viewed green product prices in 73.84 ± 11.95 trials and non-green product prices in 72.52 ± 12.71 trials per participant.

**Fig A1.**
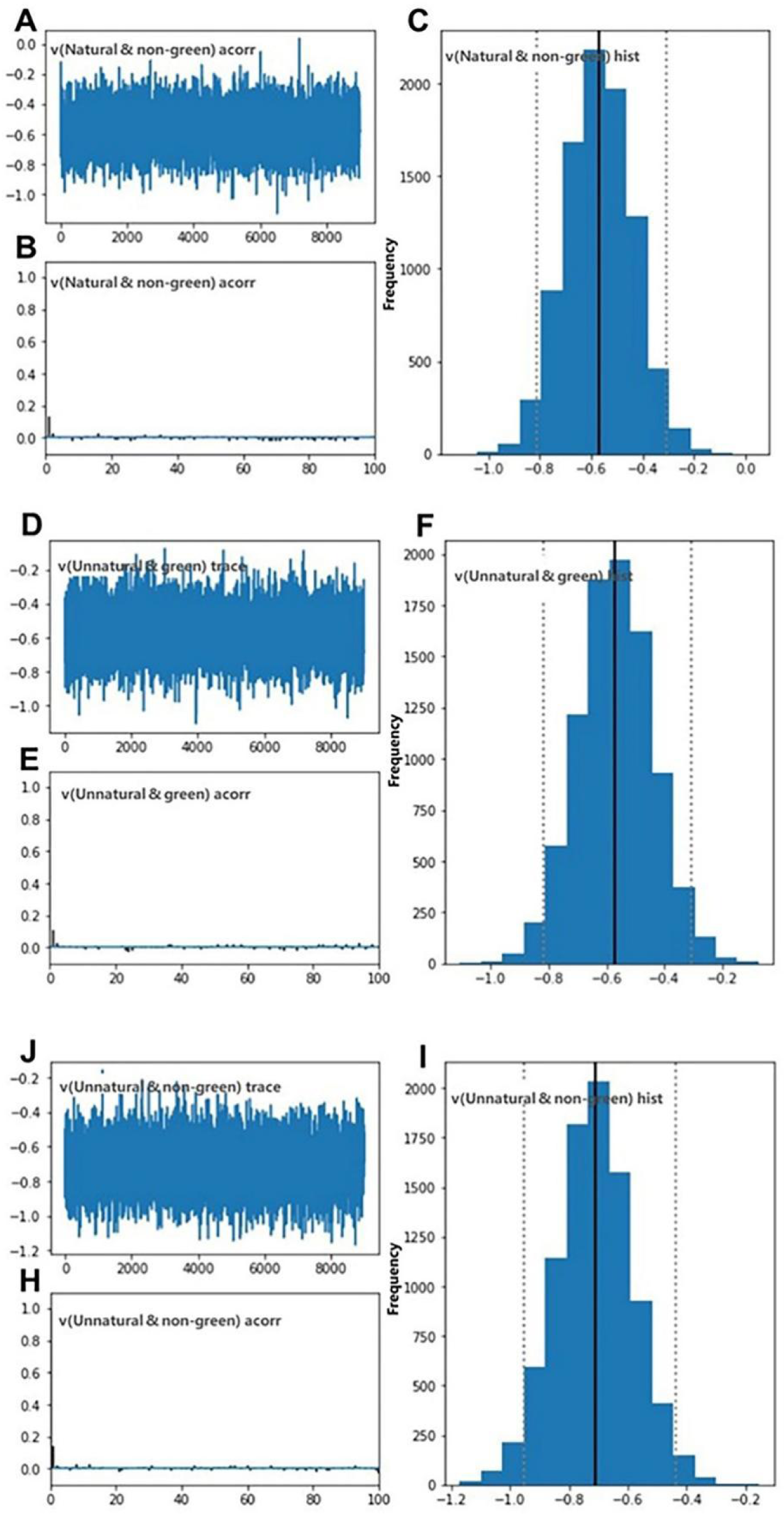
Convergence plots for the drift rate of the other three conditions. (A/D/G) The trace plot for 9000 iterations. (B/E/H) The autocorrelation for the last 100 iterations. (C/F/I) The histogram of the drift rate value.

## Footnotes

1 this label is the China Green Product Logo formulated by the China State Administration for Market Regulation in 2016; see http://www.chinagreenproduct.cn/GPIA/front.

2 Greenhouse-Geisser correction was used when the assumption of sphericity of the repeated measure was violated.

3 All pairwise comparisons used Bonferroni correction.

## Notes

### Competing Interest Statement

The authors have declared no competing interest.

## Reference

Alpert, J. I., & Alpert, M. I. (1990). Music influences on mood and purchase intentions. Psychology & Marketing, 7(2), 109–133. 10.1002/mar.4220070204

Anglada-Tort, M., Schofield, K., Trahan, T., & Müllensiefen, D. (2022). I’ve heard that brand before: the role of music recognition on consumer choice. International Journal of Advertising, 41(8), 1567–1587. 10.1080/02650487.2022.2060568

Arli, D., Tan, L. P., Tjiptono, F., & Yang, L. (2018). Exploring consumers’ purchase intention towards green products in an emerging market: The role of consumers’ perceived readiness. International Journal of Consumer Studies, 42(4), 389–401. 10.1111/ijcs.12432

Atalay, A. S., Bodur, H. O., & Rasolofoarison, D. (2012). Shining in the center: Central gaze cascade effect on product choice. Journal of Consumer Research, 39(4), 848–866. 10.1086/665984

Biswas, D. (2019). Sensory aspects of retailing: Theoretical and practical implications. Journal of Retailing, 95(4), 111–115. 10.1016/j.jretai.2019.12.001

Bradford, K. D., & Desrochers, D. M. (2009). The use of scents to influence consumers: The sense of using scents to make cents. Journal of Business Ethics, 90(2 supplement), 141–153. 10.1007/s10551-010-0377-5

Buxton, R. T., Pearson, A. L., Allou, C., Fristrup, K., & Wittemyer, G. (2021). A synthesis of health benefits of natural sounds and their distribution in national parks. Proceedings of the National Academy of Sciences, 118(14), 1–6. 10.1073/pnas.2013097118

Caldwell, C., & Hibbert, S. A. (2002). The influence of music tempo and musical preference on restaurant patrons’ behavior. Psychology & Marketing, 19(11), 895–917. 10.1002/mar.10043

Carlson, L., Grove, S. J., & Kangun, N. (1993). A content analysis of environmental advertising claims: A matrix method approach. Journal of Advertising, 22(3), 27–39. 10.1080/00913367.1993.10673409

Clement, J., Aastrup, J., & Charlotte Forsberg, S. (2015). Decisive visual saliency and consumers’ in-store decisions. Journal of Retailing and Consumer Services, 22, 187–194. 10.1016/j.jretconser.2014.09.002

Chen, J., Huang, Y., Wu, E. Q., Ip, R., & Wang, K. (2023). How does rural tourism experience affect green consumption in terms of memorable rural-based tourism experiences, connectedness to nature and environmental awareness? Journal of Hospitality and Tourism Management, 54, 166–177. 10.1016/j.jhtm.2022.12.006

Debnath, R., & Wetzel, N. (2022). Processing of task-irrelevant sounds during typical everyday activities in children. Developmental Psychobiology, 64(7), 1–17. 10.1002/dev.22331

Dong, G., Hu, Y., & Zhou, H. (2010). Event-related potential measures of the intending process: Time course and related ERP components. Behavioral and Brain Functions, 6(1), 1–7. 10.1186/1744-9081-6-15

Eurobarometer. (2022). Fairness perceptions of the green transition. Retrieved from https://europa.eu/eurobarometer/surveys/detail/2672

Evans, J. S. B. T. (2008). Dual-processing accounts of reasoning, judgment, and social cognition. Annual Review of Psychology, 59(1), 255–278. 10.1146/annurev.psych.59.103006.093629

Faul, F., Erdfelder, E., Buchner, A., & Lang, A. G. (2009). Statistical power analyses using G* Power 3.1: Tests for correlation and regression analyses. Behavior Research Methods, 41(4), 1149–1160. 10.3758/BRM.41.4.1149

Fei, M., Tan, H., Peng, X., Wang, Q., & Wang, L. (2021). Promoting or attenuating? An eye-tracking study on the role of social cues in e-commerce livestreaming. Decision Support Systems, 142, 1–10. 10.1016/j.dss.2020.113466

Ferreira, D. C. S., & Oliveira-Castro, J. M. (2011). Effects of background music on consumer behaviour: behavioural account of the consumer setting. The Service Industries Journal, 31(15), 2571–2585. 10.1080/02642069.2011.531125

Freunberger, R., Klimesch, W., Doppelmayr, M., & Höller, Y. (2007). Visual P2 component is related to theta phase-locking. Neuroscience Letters, 426(3), 181–186. 10.1016/j.neulet.2007.08.062

Frumkin, H., Bratman, G. N., Breslow, S. J., Cochran, B., Kahn Jr, P. H., Lawler, J. J., Wood, S. A. (2017). Nature contact and human health: A research agenda. Environmental Health Perspectives, 125(7), 1–18. 10.1289/EHP1663

García-de-Frutos, N., Ortega-Egea, J. M., & Martínez-del-Río, J. (2018). Anti-consumption for environmental sustainability: Conceptualization, review, and multilevel research directions. Journal of Business Ethics, 148(2), 411–435. 10.1007/s10551-016-3023-z

Gelman, A., & Rubin, D. B. (1992). Inference from iterative simulation using multiple sequences. Statistical Science, 7(4), 457–472. 10.1214/ss/1177011136

Goto, N., Mushtaq, F., Shee, D., Lim, X. L., Mortazavi, M., Watabe, M., & Schaefer, A. (2017). Neural signals of selective attention are modulated by subjective preferences and buying decisions in a virtual shopping task. Biological Psychology, 128, 11–20. 10.1016/j.biopsycho.2017.06.004

Griskevicius, V., Tybur, J. M., & Van den Bergh, B. (2010). Going green to be seen: status, reputation, and conspicuous conservation. Journal of Personality and Social Psychology, 98(3), 392–404. 10.1037/a0017346

Gruber, T., & Müller, M. M. (2005). Oscillatory brain activity dissociates between associative stimulus content in a repetition priming task in the human EEG. Cerebral Cortex, 15(1), 109–116. 10.1093/cercor/bhh113

Hayes, A. F. (2013). Introduction to Mediation, Moderation, and Conditional Process Analysis: A Regression-Based Approach. The Guilford Press.

Helmefalk, M., & Hultén, B. (2017). Multi-sensory congruent cues in designing retail store atmosphere: Effects on shoppers’ emotions and purchase behavior. Journal of Retailing and Consumer Services, 38, 1–11. 10.1016/j.jretconser.2017.04.007

Huang, Y. S., Wei, S., & Ang, T. (2021). The role of customer perceived ethicality in explaining the impact of incivility among employees on customer unethical behavior and customer citizenship behavior. Journal of Business Ethics, 178(2), 519–535. 10.1007/s10551-020-04698-9

Jacob, C. (2006). Styles of background music and consumption in a bar: an empirical evaluation. International Journal of Hospitality Management, 25(4), 716–720. 10.1016/j.ijhm.2006.01.002

Jacoby, J. (2002). Stimulus-organism-response reconsidered: an evolutionary step in modeling (consumer) behavior. Journal of consumer psychology, 12(1), 51–57. 10.1207/S15327663JCP1201_05

Jiang, H., Fan, W., & Liu, C. (2020). Event-related potential analysis on the influence of regional price and markting strategy over consumers ‘puchases decision. Revista Argentina de Clínica Psicológica, 29(1), 214–222. 10.24205/03276716.2020.28

Kaplan, S. (1995). The restorative benefits of nature: Toward an integrative framework. Journal of Environmental Psychology, 15(3), 169–182. 10.1016/0272-4944(95)90001-2

Kellert, S. R., & Wilkins, G. (1993). Dialogue with animals: Its nature and culture. In S. R. Kellert & E. O. Wilson (Eds.), The biophilia hypothesis (pp. 173–197). The Island Press.

Knutson, B., Rick, S., Wimmer, G. E., Prelec, D., & Loewenstein, G. (2007). Neural predictors of purchases. Neuron, 53(1), 147–156. 10.1016/j.neuron.2006.11.010

Koivisto, M., Jalava, E., Kuusisto, L., Railo, H., & Grassini, S. (2022). Top-down processing and nature connectedness predict psychological and physiological effects of nature. Environment and Behavior, 54(5), 917–845. 10.1177/00139165221107535

Krajbich, I., Lu, D., Camerer, C., & Rangel, A. (2012). The attentional drift-diffusion model extends to simple purchasing decisions. Frontiers in Psychology, 3, 1–18. 10.3389/fpsyg.2012.00193

Lee, E. J., Choi, H., Han, J., Kim, D. H., Ko, E., & Kim, K. H. (2020). How to “Nudge” your consumers toward sustainable fashion consumption: An fMRI investigation. Journal of Business Research, 117, 642–651. 10.1016/j.jbusres.2019.09.050

Lee, E. J., Kwon, G., Shin, H. J., Yang, S., Lee, S., & Suh, M. (2014). The spell of green: Can frontal EEG activations identify green consumers? Journal of Business Ethics, 122(3), 511–521. 10.1007/s10551-013-1775-2

Lin, Y. H. T., Hamid, N., Shepherd, D., Kantono, K., & Spence, C. (2019). Environmental sounds influence the multisensory perception of chocolate gelati. Foods, 8(4), 124. 10.3390/foods8040124

Lin, Y. H. T., Hamid, N., Shepherd, D., Kantono, K., & Spence, C. (2022a). Musical and non-musical sounds influence the flavour perception of chocolate ice cream and emotional responses. Foods, 11(12), 1784. 10.3390/foods11121784

Lin, Y. H. T., Hamid, N., Shepherd, D., Kantono, K., & Spence, C. (2022b). Sound pleasantness influences the perception of both emotional and non-emotional foods. Food Research International, 162 (PtA), 1–14. 10.1016/j.foodres.2022.111909

Linder, N. S., Uhl, G., Fliessbach, K., Trautner, P., Elger, C. E., & Weber, B. (2010). Organic labeling influences food valuation and choice. Neuroimage, 53(1), 215–220. 10.1016/j.neuroimage.2010.05.077

Lithari, C., Frantzidis, C. A., Papadelis, C. V., A. B., Klados, M. A., Kourtidou-Papadeli, C., Pappas, C., … Bamidis, P. D. (2010). Are females more responsive to emotional stimuli? A neurophysiological study across arousal and valence dimensions. Brain Topography, 23(1), 27–40. 10.1007/s10548-009-0130-5

Liu, F., Liu, S., & Jiang, G. (2022). Consumers’ decision-making process in redeeming and sharing behaviors toward app-based mobile coupons in social commerce. International Journal of Information Management, 67, 1–18. 10.1016/j.ijinfomgt.2022.102550

Liu, T., Wang, D., Wang, C., Xiao, T., & Shi, J. (2022). The influence of reward anticipation on conflict control in children and adolescents: Evidences from hierarchical drift-diffusion model and event-related potentials. Developmental Cognitive Neuroscience, 55, 1–13. 10.1016/j.dcn.2022.101118

Luck, S. J. (2005). An introduction to the event-related potential technique. The MIT Press.

Marikyan, D., Papagiannidis, S. (2023). Exercising the “Right to Repair”: A Customer’s Perspective. Journal of Business Ethics. 1–27. 10.1007/s10551-023-05569-9

Mehrabian, A., & Russell, J. A. (1974). An Approach to Environmental Psychology. The MIT Press.

Mennella, R., Vilarem, E., & Grèzes, J. (2020). Rapid approach-avoidance responses to emotional displays reflect value-based decisions: Neural evidence from an EEG study. NeuroImage, 222, 1–12. 10.1016/j.neuroimage.2020.117253

Milliman, R. E. (1982). Using background music to affect the behavior of supermarket shoppers. Journal of marketing, 46(3), 86–91. 10.2307/1251706

Michel, A., Baumann, C., & Gayer, L. (2017). Thank you for the music–or not? The effects of in-store music in service settings. Journal of Retailing and Consumer Services, 36, 21–32. 10.1016/j.jretconser.2016.12.008

Michels, N., & Hamers, P. (2023). Nature Sounds for Stress Recovery and Healthy Eating: A Lab Experiment Differentiating Water and Bird Sound. Environment and Behavior, 55(3) 175–205. 10.1177/00139165231174622

Morrin, M., & Chebat, J. C. (2005). Person-place congruency: The interactive effects of shopper style and atmospherics on consumer expenditures. Journal of Service Research, 8(2), 181–191. 10.1177/10946705052794

North, A. C., & Hargreaves, D. J. (1998). The effect of music on atmosphere and purchase intentions in a cafeteria. Journal of Applied Social Psychology, 28(24), 2254–2273. 10.1111/j.1559-1816.1998.tb01370.x

North, A. C., Sheridan, L. P., & Areni, C. S. (2016). Music congruity effects on product memory, perception, and choice. Journal of Retailing, 92(1), 83–95. 10.1016/j.jretai.2015.06.001

North, A. C., Shilcock, A., & Hargreaves, D. J. (2003). The effect of musical style on restaurant customers’ spending. Environment and Behavior, 35(5), 712–718. 10.1177/0013916503254749

Oakes, S. (2000). The influence of the musicscape within service environments. Journal of services marketing, 14(7), 539–556. 10.1108/08876040010352673

Ozkara, B. Y., & Bagozzi, R. (2021). The use of event related potentials brain methods in the study of conscious and unconscious consumer decision making processes. Journal of Retailing and Consumer Services, 58, 1–17. 10.1016/j.jretconser.2020.102202

Peng-Li, D., Andersen, T., Finlayson, G., Byrne, D. V., & Wang, Q. J. (2022). The impact of environmental sounds on food reward. Physiology & Behavior, 245, 1–10. 10.1016/j.physbeh.2021.113689

Peng-Li, D., Mathiesen, S. L., Chan, R. C., Byrne, D. V., & Wang, Q. J. (2021). Sounds Healthy: Modelling sound-evoked consumer food choice through visual attention. Appetite, 164, 1–17. 10.1016/j.appet.2021.105264

Pisauro, M. A., Fouragnan, E., Retzler, C., & Philiastides, M. G. (2017). Neural correlates of evidence accumulation during value-based decisions revealed via simultaneous EEG-fMRI. Nature Communications, 8(1), 1–9. 10.1038/ncomms15808

Ratcliff, R., Smith, P. L., Brown, S. D., & McKoon. (2016). Diffusion Decision Model: Current Issues and History. Trends in Cognitive Sciences, 20(4), 260–281. 10.1016/j.tics.2016.01.007

Ratcliff, R., & Rouder, J. N. (1998). Modeling response times for two-choice decisions. Psychological Science, 9(5), 347–356. 10.1111/1467-9280.00067

Ratcliffe, E., Gatersleben, B., & Sowden, P. T. (2013). Bird sounds and their contributions to perceived attention restoration and stress recovery. Journal of Environmental Psychology, 36, 221–228. 10.1016/j.jenvp.2013.08.004

Regenbogen, C., Johansson, E., Andersson, P., Olsson, M. J., & Lundstrom, J. N. (2016). Bayesian-based integration of multisensory naturalistic perithreshold stimuli. Neuropsychologia, 88, 123–130. 10.1016/j.neuropsychologia.2015.12.017

Roberts, I. D., & Hutcherson, C. A. (2019). Affect and decision making: Insights and predictions from computational models. Trends in Cognitive Sciences, 23(7), 602–614. 10.1016/j.tics.2019.04.005

Robertson, D. C., Voegtlin, C., & Maak, T. (2017). Business ethics: The promise of neuroscience. Journal of Business Ethics, 144(4), 679–697. 10.1007/s10551-016-3312-6

Schwartz, D., Loewenstein, G., & Agüero-Gaete, L. (2020). Encouraging pro-environmental behaviour through green identity labelling. Nature Sustainability, 3(9), 746–752. 10.1038/s41893-020-0543-4

Shang, Q., Jin, J., Pei, G., Wang, C., Wang, X., & Qiu, J. (2020). Low-order webpage layout in online shopping facilitates purchase decisions: evidence from event-related potentials. Psychology Research and Behavior Management, 13, 29–39. 10.2147/PRBM.S238581

Smith, P. C., & Curnow, R. (1966). “ Arousal hypothesis” and the effects of music on purchasing behavior. Journal of applied psychology, 50(3), 255–256. 10.1037/h0023326

Smith, P. L., & Ratcliff, R. (2004). Psychology and neurobiology of simple decisions. Trends in neurosciences, 27(3), 161–168. 10.1016/j.tins.2004.01.006

Sønderskov, K. M., & Daugbjerg, C. (2011). The state and consumer confidence in eco-labeling: organic labeling in Denmark, Sweden, The United Kingdom and The United States. Agriculture and Human Values, 28, 507–517. 10.1007/s10460-010-9295-5

Spendrup, S., Hunter, E., & Isgren, E. (2016). Exploring the relationship between nature sounds, connectedness to nature, mood and willingness to buy sustainable food: a retail field experiment. Appetite, 100, 133–141. 10.1016/j.appet.2016.02.007

Stanton, S. J., Sinnott-Armstrong, W., & Huettel, S. A. (2017). Neuromarketing: Ethical implications of its use and potential misuse. Journal of Business Ethics, 144(4), 799–811. 10.1007/s10551-016-3059-0

Sun, L., Zhao, Y., & Ling, B. (2020). The joint influence of online rating and product price on purchase decision: An EEG study. Psychology Research and Behavior Management, 13, 291–301. 10.2147/PRBM.S238063

Tandon, A., Sithipolvanichgul, J., Asmi, F., Anwar, M. A., & Dhir, A. (2023). Drivers of green apparel consumption: Digging a little deeper into green apparel buying intentions. Business Strategy and the Environment, 32(6), 3997–4012. 10.1002/bse.3350

Telpaz, A., Webb, R., & Levy, D. J. (2015). Using EEG to predict consumers’ future choices. Journal of marketing research, 52(4), 511–529. 10.1509/jmr.13.0564

Van Hooff, J. C., Crawford, H., & Van Vugt, M. (2011). The wandering mind of men: ERP evidence for gender differences in attention bias towards attractive opposite sex faces. Social cognitive and affective neuroscience, 6(4), 477–485. 10.1093/scan/nsq066

Wang, Y., Yang, X., Tang, Z., Xiao, S., & Hewig, J. (2021). Hierarchical neural prediction of interpersonal trust. Neuroscience Bulletin, 37, 511–522. 10.1007/s12264-021-00628-5

Wiecki, T. V., Sofer, I., & Frank, M. J. (2013). HDDM: Hierarchical Bayesian estimation of the drift-diffusion model in Python. Frontiers in neuroinformatics, 14, 1–10. 10.3389/fninf.2013.00014

Xie, Z., Yu, Y., Zhang, J., & Chen, M. (2022). The searching artificial intelligence: Consumers show less aversion to algorithm-recommended search product. Psychology & Marketing, 39(10), 1902–1919. 10.1002/mar.21706

Xu, Y., Hamid, N., Shepherd, D., Kantono, K., Reay, S., Martinez, G., & Spence, C. (2019). Background soundscapes influence the perception of ice-cream as indexed by electrophysiological measures. Food Research International, 125, 1–11. 10.1016/j.foodres.2019.108564

Lin, Y. H. T., Hamid, N., Shepherd, D., Kantono, K., & Spence, C. (2022). Musical and non-musical sounds influence the flavour perception of chocolate ice cream and emotional responses. Foods, 11(12), 1–21. 10.3390/foods11121784

Yan, L., Keh, H. T., & Chen, J. (2021). Assimilating and differentiating: the curvilinear effect of social class on green consumption. Journal of Consumer Research, 47(6), 914–936. 10.1093/jcr/ucaa041

Yun, J. H., Kim, Y., & Lee, E. J. (2022). ERP study of liberals’ and conservatives’ moral reasoning processes: Evidence from South Korea. Journal of Business Ethics, 176(4), 723–739. 10.1007/s10551-021-04734-2

Zaremohzzabieh, Z., Ismail, N., Ahrari, S., & Samah, A. A. (2021). The effects of consumer attitude on green purchase intention: A meta-analytic path analysis. Journal of Business Research, 132, 732–743. 10.1016/j.jbusres.2020.10.053

Zhang, J., Yun, J. H., & Lee, E. J. (2021). Brain buzz for Facebook? Neural indicators of SNS content engagement. Journal of Business Research, 130, 444–452. 10.1016/j.jbusres.2020.01.029

Zhang, S., Ren, S., & Tang, G. (2023). From Passive to Active: The Positive Spillover of Required Employee Green Behavior on Green Advocacy. Journal of Business Ethics, 1–20. 10.1007/s10551-023-05494-x

Zheng, X., Men, J., Yang, F., & Gong, X. (2019). Understanding impulse buying in mobile commerce: An investigation into hedonic and utilitarian browsing. International Journal of Information Management, 48, 151–160. 10.1016/j.ijinfomgt.2019.02.010

Zioga, I., Harrison, P. M., Pearce, M. T., Bhattacharya, J., & Luft, C. D. B. (2020). From learning to creativity: Identifying the behavioural and neural correlates of learning to predict human judgements of musical creativity. Neuroimage, 206, 1–19. 10.1016/j.neuroimage.2019.116311

